# Phrank measures phenotype sets similarity to greatly improve Mendelian diagnostic disease prioritization

**DOI:** 10.1101/225854

**Authors:** Karthik A. Jagadeesh, Johannes Birgmeier, Harendra Guturu, Cole A. Deisseroth, Aaron M. Wenger, Jonathan A. Bernstein, Gill Bejerano

## Abstract

**Purpose:** Exome sequencing and diagnosis is beginning to spread across the medical establishment. The most time-consuming part of genome based diagnosis is the manual step of matching the potentially long list of patient candidate genes to patient phenotypes to identify the causative disease.

**Methods:** We introduce Phrank (for phenotype ranking), an information-theory inspired method that utilizes a Bayesian Network to prioritize candidate diseases or genes, as a stand-alone module that can be run with any underlying knowledgebase and any variant filtering scheme.

**Results:** Phrank outperforms existing methods at ranking the causative disease or gene when applied to 169 real patient exomes with Mendelian diagnoses. Phrank’s greatest improvement is in disease space, where across all 169 patients it ranks only 3 diseases on average ahead of the true diagnosis, whereas Phenomizer ranks 32 diseases ahead of the causal one.

**Conclusion:** Using Phrank to rank all patient candidate genes or diseases, as they start working through a new case, will save the busy clinician much time in deriving a genetic diagnosis.

## Main Text

### Introduction

Genomic data is becoming increasingly utilized in medical genetics clinical practice^1–3^. Exome sequencing has allowed for the identification of the genetic basis of thousands of different Mendelian disorders^4–6^. In most clinical sequencing cases, only the proband’s exome is sequenced. Such a typical “singleton” patient exome contains 100-300 variants of uncertain significance (VUS), of which only one or two may adversely affect a single gene of interest.

Clinicians ultimately diagnose patients by identifying the disease that best explains the patient’s phenotypes^7^. To do this, they mentally calculate a “fuzzy” match between the patient phenotype and all diseases they are aware of to try and find a disease that best fits the phenotypes observed in the patient. Clinicians also try to read up on the different patient candidate genes, to see whether any gene has been previously implicated in causing phenotypes similar to those in their patient. This is no easy task, with several thousands of well-characterized Mendelian diseases, caused by well over 3,000 known Mendelian disease genes^8–10^, and hundreds of novel gene-disease associations discovered annually^7^. On average, clinicians analyzing a patient with a Mendelian disorder have estimated spending 54 minutes to examine a single variant’s pathogenicity^11^. In some cases clinicians can immediately recognize the genetic basis of the disease, but in general, identifying a patient’s causative gene has been estimated to consume on average a workweek of expert time^12–14^.

With 7 million (5.4%) of babies born each year worldwide with serious inherited genetic disorders^15^, and genetic testing workflows capable of sequencing and generating variant call files for hundreds of individuals per day, busy clinicians are in great need of computational approaches to aid them in making a rapid accurate diagnosis. Such automated methods attempt to rank all patient possibly-pathogenic variants for their known ability to explain the patient set of phenotypes. The goal is to bring the causal gene to the top of the patient ranked list, such that when the busy clinician first lays eyes on a sequenced case, their attention is drawn to the most likely diagnostic hypotheses as early as possible. Different computational tools attempt to achieve this goal by combining computable measures of similarity between a patient set of phenotypes and the set of phenotypes associated with any gene on the patient list, observed variant frequency in the general population and measures of predicted variant pathogenicity^16–19^.

Most automated phenotype similarity based ranking methods such as Phevor^20^ and PhenIX^21^ rank genes, and leave the clinician with the final step of identifying the causative disease.

Phenomizer^22^ is one of few disease ranking tools, making it popular among clinicians. We introduce a novel information theory inspired ranking method, Phrank, which greatly improves on Phenomizer for identifying the causative disease. For maximum utility, Phrank is offered both through the AMELIE web interface^23^ where it is deployed with a particular knowledgebase, as well as in the form of a code module that can be combined with any variant ranking scheme and any underlying knowledgebase.

### Materials and Methods

Patient variant prioritization is ultimately determined via a combination of multiple variant and host gene properties. Phrank isolates and optimizes one important feature: scoring each disease caused by a patient candidate gene for its ability to explain the set of patient phenotypes.

#### General Phrank inputs: phenotype ontology and a gene-disease-phenotype knowledgebase

Phrank assumes access to a phenotype ontology representing relationships between phenotypes as a rooted directed acyclic graph (DAG), such as the Human Phenotype Ontology^10^ (HPO) and a knowledgebase of gene-disease-phenotype relationships, such as the Human Phenotype Ontology Annotations (HPO-A), where each entry consists of a diagnostic gene, a disease caused by mutations in the gene, and a disease associated phenotype. By definition, whenever a gene *g* is annotated to cause phenotype *φ*, it is considered to also cause (a particular instance of) all phenotypes that are ancestral to *φ*, denoted as the set *anc* (*φ*). For example, if *φ* is elbow hypertrichosis, *anc*(*φ*) will contain elbow hypertrichosis itself, as well as hypertrichosis, abnormal hair quantity, abnormality of the hair, etc. up to the root of the phenotype DAG. For a set of phenotypes Φ, *anc*(Φ) is the union of *anc*(*φ*_*i*_) for every *φ*_*i*_ in Φ.

#### Computing the conditional probability of a phenotype in a DAG

Using the DAG and knowledgebase, we define |*G*_φ_| to be the number of genes associated with phenotype φ, and |*G*_*pa(φ)*_| to be the number of genes associated with any parent phenotype of φ in the graph. We overlay a Boolean Bayesian Network^24^ on the HPO DAG. In a Bayesian Network, each node is assigned a probability of being observed conditioned on its parent nodes being observed. Here, we define *P(root)* = 1, and the conditional probability of observing phenotype *φ* given its parent phenotypes 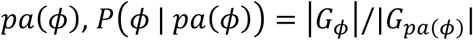 if all parent phenotypes *pa(φ)* are observed and let the conditional probability 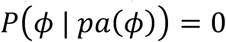 otherwise.

#### Phrank ranks candidate genes and diseases using a novel phenotype similarity measurement

Per patient, Phrank takes as input a list of patient phenotypes and a list of patient candidate genes. Based on user preference, Phrank will then output a score for each gene in the candidate gene list, or for each disease that according to the knowledgebase may be caused by one or more of the genes on the patient candidate gene list. The higher the score, the better the gene or disease is thought to explain the provided set of patient phenotypes.

Phrank computes a score measuring the similarity between a patient’s phenotypes and each gene/disease-phenotype set. Each phenotype’s contribution to the Phrank score is a function of the number of genes known to cause the phenotype. Intuitively, the fewer genes are known to cause a phenotype, the more impressive is a candidate gene’s ability to explain this patient phenotype, and the higher the score derived from such a match (see Figure 1 and below for details).

**Figure 1.**
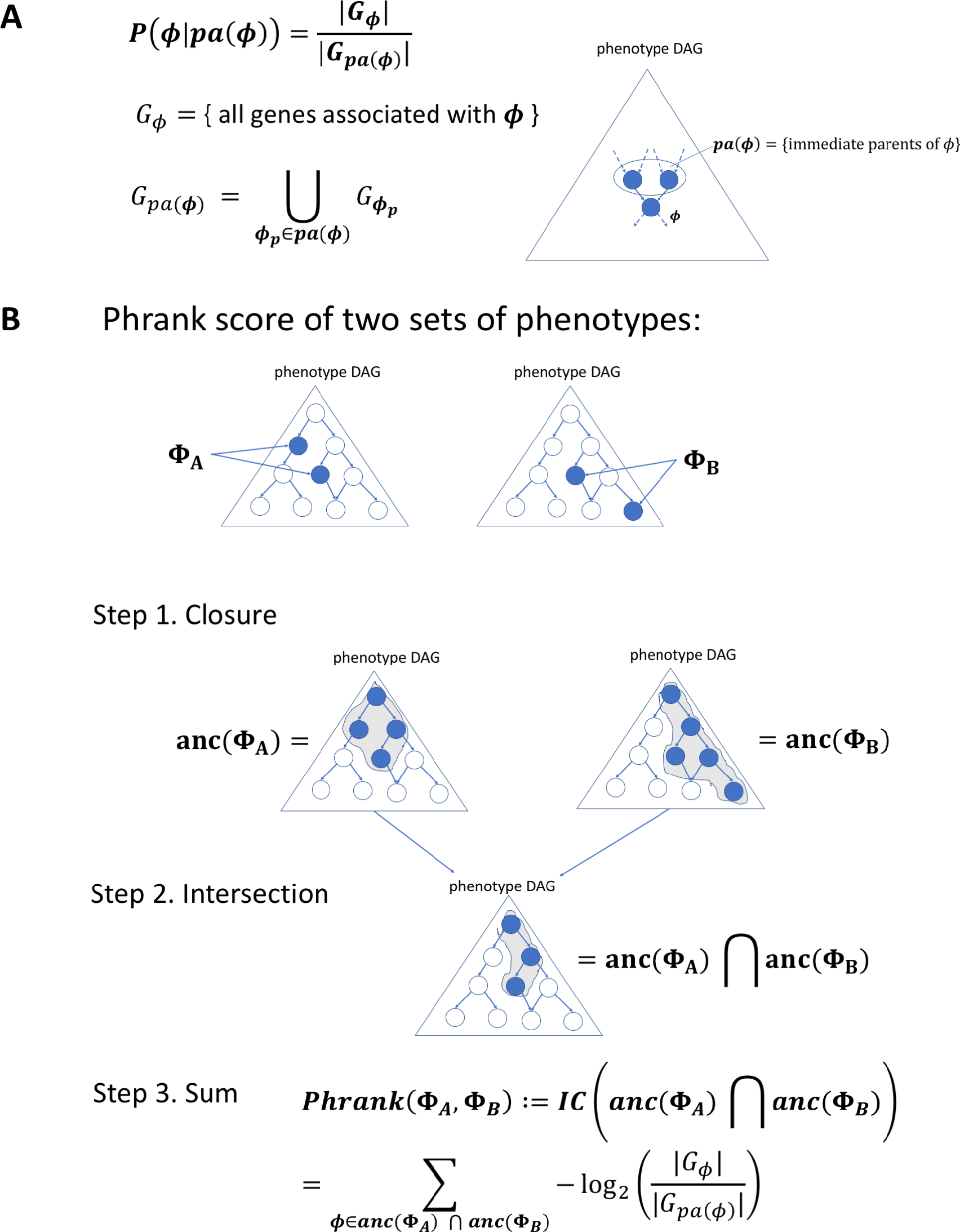
The Phrank information content based score. (A) The conditional probability *P(φ|pa(φ))* for a single node in the phenotype ontology. As these are log transformed, negated and then summed in panel B via paths to the root node, one can see that nodes annotated with fewer genes will ultimately contribute more to the final score (as the match they represent is less expected by chance; see Methods). (B) The Phrank match score of any two sets of phenotypes, Φ_A_ and Φ_B_, is defined as the information content of the intersection of all ancestral nodes of both sets, *anc*(*Φ*_A_) and *anc*(*Φ*_B_) and can be computed as shown in Step 3 (see Supplemental Note 1 for the derivation). We define the Phrank score of any disease to any particular patient as the Phrank score of the set of phenotypes associated with the disease and the set of phenotypes observed in the patient (see Methods).

#### Defining a Phrank score between any two sets of phenotypes

Given two sets of phenotypes, Φ_*A*_ and Φ_*B*_, we define the Phrank score between them as the information content of the intersection of the ancestral closure of the two sets (see Figure 1). In Supplemental Note 1 we show that this quantity can be computed using

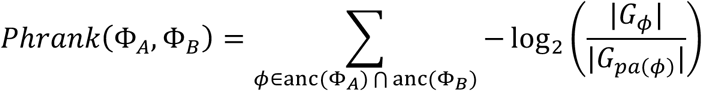

#### The (patient-specific) Phrank score of genes and diseases

Given the above definition of the Phrank score between any two sets of phenotypes, the patient-specific <u>Phrank score of a disease</u> is simply the Phrank score between the set of phenotypes associated with the disease in our knowledgebase and the input set of patient phenotypes. The <u>Phrank score of a gene</u> is defined as the maximal Phrank score for a disease that can be caused by the gene, according to the knowledgebase.

#### Human Phenotype Ontology (HPO) and HPO Annotations (HPO-A)

To test Phrank’s performance we use the popular Human Phenotype Ontology^10^ (HPO). HPO is a rooted directed acyclic graph (DAG) that includes a hierarchical comprehensive description of human phenotypes. The phenotype DAG root node is labeled “Phenotypic Abnormality” (HP:0000118). Parent-child “is a” relationships that are part of the graph consist of a more general parent term and a more specific child term. For example, the term “Hypertrichosis” is the parent of the term “Elbow hypertrichosis”. A term can have multiple children, and multiple parents.

The Human Phenotype Ontology project also curates gene-phenotype-disease relationships^10^ based on the Online Mendelian Inheritance in Man (OMIM)^8^ manually curated knowledgebase. We refer to these as HPO-A (HPO Annotations), and have downloaded them from http://compbio.charite.de/jenkins/job/hpo.annotations.monthly/. HPO build 127 provides a rooted DAG over 11,156 distinct phenotypes, and HPO-A provides direct mappings between 3,406 genes, 3,995 diseases, and 5,640 HPO phenotypes.

#### Benchmark set of 169 real diagnosed patients

A dataset of diagnosed patients was downloaded from the Deciphering Developmental Disorders (DDD) study^25^ in the European Genome-Phenome Archive (EGA)^26^ study EGAS00001000775. The DDD study recruited patients satisfying a wide range of neurodevelopmental disorders and congenital anomalies^26^. The dataset contains patient Variant Call Files (VCFs), a list of HPO phenotypes and the causative gene for each patient. From this set we removed cases with a sibling already in the set, cases offering a novel causative gene hypothesis and cases where the disease was caused by large structural or mosaic variants. A board-certified clinical geneticist on our team independently reviewed these patients (without the use of a gene- or disease-ranking tool) and identified OMIM disease IDs that best explain each patient’s condition and causal gene. A total of 169 patients suffering from nearly a hundred different diseases were found to have high confidence diagnoses (see Supplementary Table 1).

#### Variant annotation

ExAC^13^ v0.3 and the 1000 Genomes Project^12^ (KGP) were used to annotate variants with observed control population frequencies. ANNOVAR^28^ v527 was used to annotate variants with predicted effect on protein coding genes using gene isoforms from the ENSEMBL gene set^29,30^ version 75 for the hg19/GRCh37 assembly of the human genome.

#### Variant filtering

We observed on average a total of 98,815 variants per individual. Patient variants are filtered to only keep those predicted by ANNOVAR to be nonsynonymous, stopgain, stoploss, splice affecting, frameshift indel or nonframeshift indel mutations^7^. Further, variants are filtered to a set of candidate disease causing mutations based on allele frequency in the control population. Genetic variants are filtered to only keep those with an allele frequency of 0.1% or less in any ExAC or KGP population if they are heterozygous and do not co-occur with at least one other variant in the same transcript. Similarly, variants are filtered to only keep those with an allele frequency of 0.5% or less if they are homozygous or co-occur with at least one other variant in the same transcript. There are on average 281 variants per individual after following this common filtration strategy^31^. Variants are considered to be likely benign if they do not satisfy these criteria. The causative variant in all 169 patients satisfy these criteria and are properly retained in the respective candidate variant/gene list.

#### Phenomizer, Phevor and PhenIX gene/disease rankings

Phenomizer^22^ disease similarity scores were obtained for each patient by entering the patient’s HPO phenotypes into the Phenomizer website (http://compbio.charite.de/phenomizer/) and subsetting the output ranked list of all diseases to just those associated with a patient’s candidate genes containing possibly pathogenic variants.

Gene rankings by Phevor^20^ were obtained for each patient by entering the patient’s HPO phenotypes into Phevor’s website (http://weatherby.genetics.utah.edu/phevor2/index.html) using an input file containing all 20,745 protein-coding genes in Ensembl build 75 and then subsetting the returned ranked list of all input protein-coding genes to the list of patient’s candidate genes.

Gene rankings by PhenIX^21^ were obtained for each patient by running Exomiser^32^, an integrated variant filtering and prioritization tool, with no filters, on each patient’s filtered VCF file containing the patient’s possibly pathogenic variants.

#### Using AMELIE as the underlying knowledgebase

AMELIE^23^ (for <u>A</u>utomatic <u>M</u>endelian <u>L</u>iterature <u>E</u>valuation) is an alternative Mendelian gene – phenotype association knowledgebase. It is populated entirely using a Natural Language Processing and Machine Learning approach, directly from the full text primary literature itself^23^. AMELIE links 12,295 genes to 6,669 HPO phenotypes. AMELIE does not extract disease names. Instead, it effectively trades the notion of a disease with that of a full text paper. Every causal gene it extracts from a paper is linked to a set of HPO phenotypes extracted with it from the same paper. In order to compare Phrank on HPO-A to Phrank on AMELIE, we used the Phrank HPO-A based gene ranking as above. For AMELIE, paper replaced disease. In other words, the patient-specific <u>AMELIE-based Phrank score of a gene</u> was defined as the maximal Phrank score of any paper about the gene according to the AMELIE knowledgebase.

#### Phrank Code Availability

The Phrank source code will be available at https://bitbucket.org/bejerano/phrank.

### Results

#### Patient benchmark set

Previous methods largely used simulated patients with artificially assigned phenotypes for performance evaluation^20–22^. On these synthetic sets, some of the methods we compare to ranked the causative gene at the top in 90% of cases. Taking advantage of the growing availability of real patient data, we curated a set of 169 patients from the Deciphering Developmental Diseases (DDD) study^25^, as described above (Methods and Supplementary Table 1). On average, each patient in the final set is characterized by 7.5 phenotypes, and carries a candidate list of 278.8 genes. As we show next, this real data set poses a much bigger challenge to all tested methods.

#### Phrank greatly improves disease ranking

We gave Phrank as input each patient candidate gene list, along with their respective set of phenotypes. Phenomizer^22^ only takes as input the set of patient phenotypes, to rank all possible diseases. Both Phrank and Phenomizer provided as output a list of diseases ranked for their ability to explain the patient set of phenotypes. In order to compare the two methods, the Phenomizer output was subset only to diseases that according to OMIM can be caused by genes on the patient candidate list. For both methods, the rank of the correct disease diagnosis was noted. Phenomizer ranked the causative disease at the top in 3.6% of patients, in the top 5 in 17.2% of patients and in the top 10 in 31.4% of patients. Across all 169 cases, Phenomizer ranks an average of 32 diseases ahead of the patient’s actual disease. In comparison, Phrank ranked the causative disease at the top in 26% of patients, in the top 5 in 55% of patients and in the top 10 in 64.5% of patients (Figure 2A), outperforming Phenomizer at all ranking thresholds. On average, Phrank ranks only 3 diseases before the patient’s actual disease. Phrank significantly outperforms Phenomizer (p≤2.2*10^−16^, Wilcoxon signed rank test). Assuming equal time to evaluate each disease hypothesis, Phenomizer reduces clinician average time spent per patient by 25% compared to a randomly shuffled disease list baseline while Phrank reduces it by 90.9%, thus greatly accelerating diagnosis.

**Figure 2.**
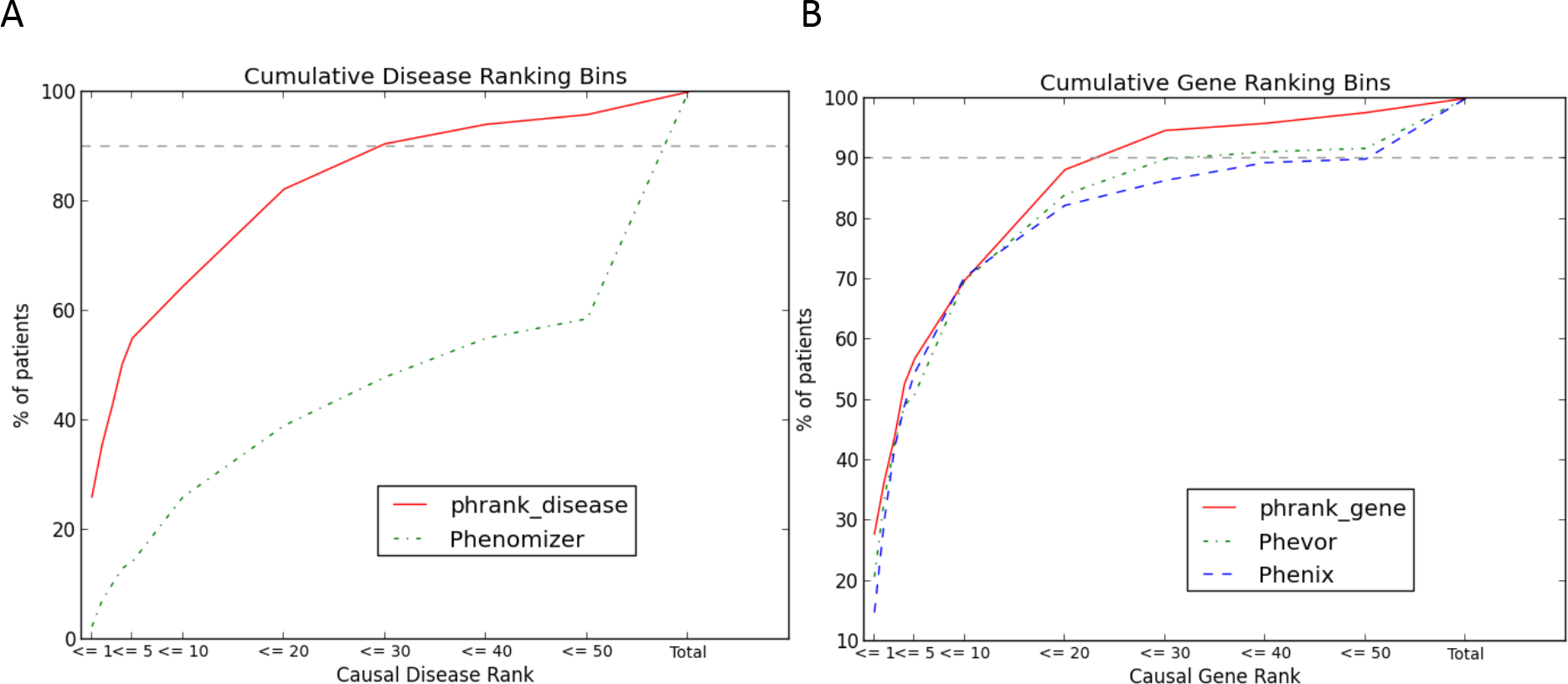
Phrank performance on a set of 169 diagnosed patients. A) The fraction of cases correctly diagnosed (y-axis) is plotted against a cumulative causal disease rank (x-axis: is the correct disease diagnosis ranked first? Is it ranked in the top 5? etc.) for both Phenomizer and Phrank. Phrank outperfoms Phenomizer significantly (p≤2.2*10^−16^, Wilcoxon signed rank test) as Phrank ranks the causative disease higher than Phenomizer at all ranking thresholds. B) The fraction of cases correctly diagnosed (y-axis) is plotted against a cumulative causal gene rank (x-axis: is the correct gene diagnosis ranked first? Is it ranked in the top 5? etc.) for Phevor, PhenIX and Phrank. Phrank slightly outperforms PhenIX and Phevor (p≤0.19 and p≤0.41, Wilcoxon signed rank test, respectively), but the difference is not statistically significant. Over all cases Phrank gives the causative gene an average rank of 9.5, while PhenIX and Phevor achieve average ranks of 15.5 and 21, respectively.

#### Phrank modestly improves gene ranking

While ranking diseases enables the most natural way to convey recommendations to the attending clinician, some tools like Phevor^20^ and PhenIX^21^, rank genes and not diseases. The Phrank score can be easily converted to this task by assigning to each gene the score of the highest scoring disease it is known to cause (Figure 1 and Methods). Phrank, PhenIX and Phevor results for each patient set of phenotypes was subset to the patient candidate gene list, and the rank of each known causative gene was collected. PhenIX and Phevor ranked the causative gene first in 15.4% and 21.3% of patients, in the top 5 in 54.4% and 52.7% of patients, respectively. In comparison, Phrank ranked the causative gene at the top in 27.8% of patients, and in the top 5 in 56.8% of patients (Figure 2B). Over all cases Phrank gives the causative gene an average rank of 9.5, while Phevor and PhenIX only achieve an average rank of 21 and 15.5, respectively.

#### Phrank works with any underlying knowledgebase

Some existing tools combine their ranking algorithm and underlying knowledgebase such that tool users can only use the algorithm with the provided knowledgebase. Phrank completely decouples the two, and can be used with any appropriately populated knowledgebase. For example, HPO-A^10^ is mostly based on manually curated gene-disease-phenotype associations, whereas AMELIE^23^ uses Machine Learning to extract Mendelian gene – phenotype associations directly from the primary literature itself (see Methods). HPO-A and AMELIE have been compared elsewhere^23^. Our goal here is only to show that Phrank can easily be used interchangeably with both knowledgebases to reveal pros and cons of each approach as they reflect in their ranking of the different cases. In particular, using Phrank, AMELIE outperforms HPO-A at all thresholds over our patient set (Figure 3).

**Figure 3.**
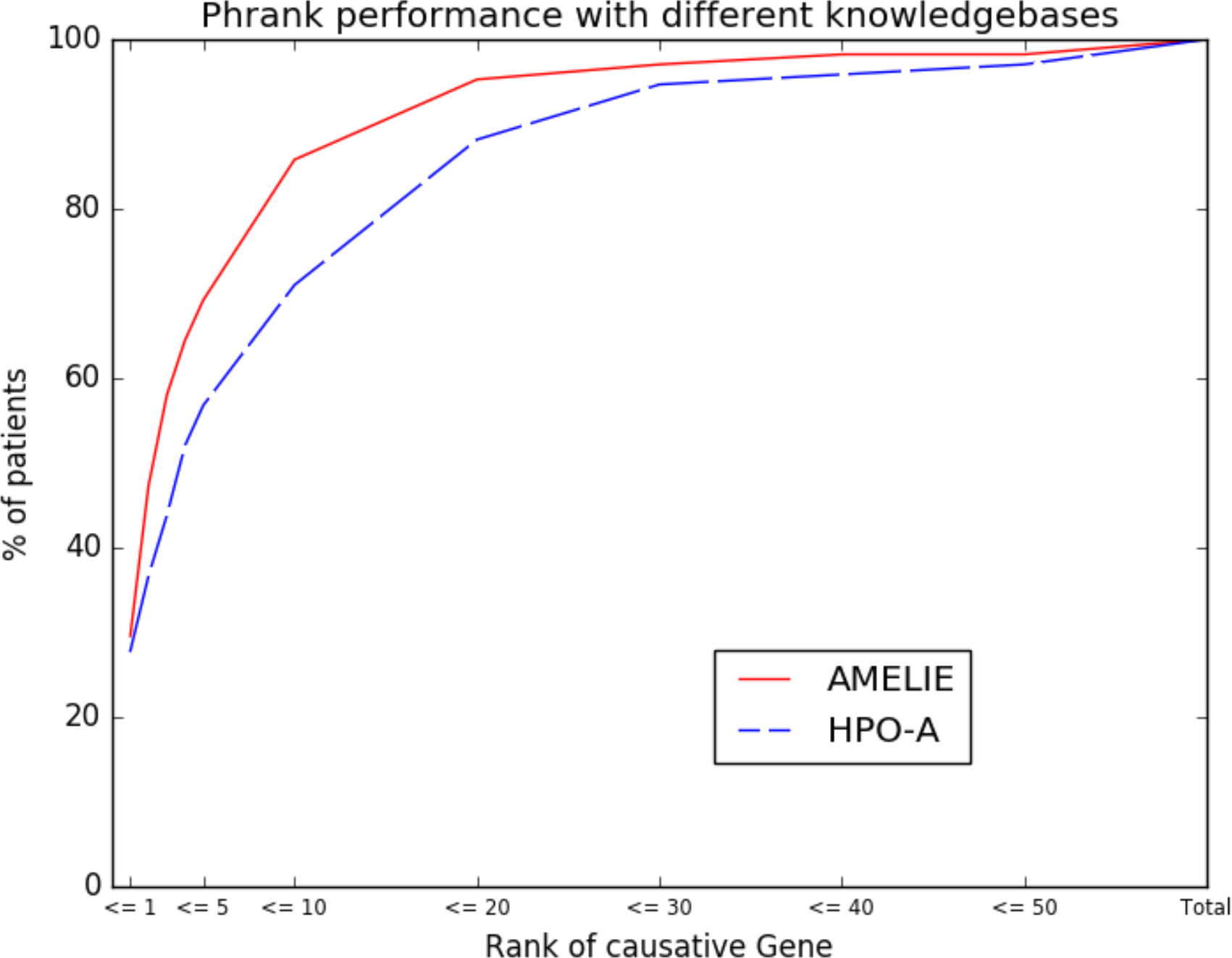
Phrank can be easily used with different knowledgebases. We show the performance of the Phrank algorithm when using the Human Phenotype Ontology^10^ annotations (HPO-A) knowledgebase based on manually curated relationships, and the AMELIE^23^ knowledgebase curated using natural language processing (NLP) from the primary literature.

### Discussion

Phrank aspires to enable clinicians to accelerate diagnosis of Mendelian diseases by ranking more appealing disease or gene matches to each patient higher. Here, we compare Phrank to Phenomizer, a disease ranking method, and to Phevor and PhenIX, two gene ranking methods. Phrank uses a natural information theoretic formulation that quantifies how informative a phenotype is for diagnosis to measure phenotype sets similarity. It uses the number of genes known to cause a phenotype as a proxy for the likelihood of observing the phenotype itself.

Computing *anc*(Φ) paired with the conditional probabilities allows neighboring phenotypes (e.g., from imprecise/over precise phenotype annotation) to be included in the final similarity score without double counting the contribution from shared ancestors.

All three methods we compare to were previously benchmarked on simulated patients, likely due to the paucity of real patient cases. To evaluate these existing methods, phenotypes for simulated patients were drawn directly from gene-phenotype knowledgebases with some noise perturbation. The same knowledgebases are used by the algorithm for gene or disease ranking, likely resulting in somewhat overly optimistic performance. Indeed, on simulated data, Phenomizer reported that the causative disease is ranked at the top in over 75% of patients^22^ and Phevor^20^ and PhenIX^21^ reported that the causative gene is ranked at the top in over 90% of patients. As shown in our study (Figure 2), when using these phenotype-based ranking methods on real patients with clinician-noted phenotypes, performance drops significantly. We recommend that future disease- and gene-ranking methods use real patient sets, such as our supplementary Table 1 EGA set, or other, to test their approaches.

Phrank is the first published method we are aware of that explicitly decouples its ranking method from both the underlying knowledgebase and the variant filtering scheme. Phrank is offered via both the ready to use AMELIE^23^ portal, as well as via a simple to use code package, offering clinicians maximum flexibility in incorporating Phrank into their preferred workflow.

We have focused our comparisons to Phevor, PhenIX and Phenomizer because these methods for disease and gene ranking use minimal data beyond the phenotype DAG and annotations. Other tools such as Phenolyzer^33^, eXtasy^34^ and Exomiser^32^ implement full variant prioritization methods incorporating a phenotype similarity measure, frequency based variant filters, pathogenicity scores and other helpful information. Phrank can effectively be used as the phenotype similarity measure and be incorporated into any such variant prioritization method.

While Phrank improves gene- and particularly disease-ranking methods, our set of real patients makes it clear there is still much room for improvement. Ideally, the causative gene or disease should be top 5 if not at the top of the ranked list of candidates with over 90% confidence. This improvement may come from improvements to the phenotype similarity algorithm, the underlying phenotype ontology’s relationship structure and/or the knowledgebase of gene-disease-phenotype associations. Such improvements will be critical in handling the ever growing flow of genomic data produced in the quest to better patient lives.

## Acknowledgements

We thank Yosuke Tanigawa, Ethan Dyer, Golan Yona and all other members of the Bejerano lab for valuable discussions and project feedback. We would also like to thank the European Genome-Phenome Archive (EGA) and the Deciphering Developmental Diseases (DDD) project. The DDD study presents independent research commissioned by the Health Innovation Challenge Fund [grant number HICF-1009-003], a parallel funding partnership between the Wellcome Trust and the Department of Health, and the Wellcome Trust Sanger Institute [grant number WT098051]. The views expressed in this publication are those of the author(s) and not necessarily those of the Wellcome Trust or the Department of Health. The study has UK Research Ethics Committee approval (10/H0305/83, granted by the Cambridge South REC, and GEN/284/12 granted by the Republic of Ireland REC). The research team acknowledges the support of the National Institute for Health Research, through the Comprehensive Clinical Research Network. as well as the patients and professionals involved in the Deciphering Developmental Disorders (DDD) study deposited in the European Genome Archive (EGA). This work was funded in part by the Stanford Graduate Fellowship and CEHG Fellowship to K.A.J., a Bio-X Stanford Interdisciplinary Graduate Fellowship to J.B., the Stanford Pediatrics Department, DARPA, a Packard Foundation Fellowship and a Microsoft Faculty Fellowship to G.B.

## Author Contributions

K.A.J., J.B., H.G. and A.M.W. wrote software tools used for the analysis. K.A.J., J.B., C.A.D. and G.B. ran the experiments and analyzed results. J.A.B. provided detailed clinician analysis of the patient dataset. K.A.J., J.B. and G.B. wrote the manuscript. G.B. supervised the study.

---

**Supplementary Table 1. Patient set**. A set of 169 DDD patients, the number of genes containing variants of unknown significance (VUS), the causal gene and causative disease in each.

## Supplementary Note 1

### Computing the Phrank score

We overlay a Bayesian Network^24^ on the HPO DAG. A Bayesian Network is a DAG in which every node is associated with a conditional probability distribution dependent only on its parents. Let *NonDesc*_*φ*_ be the set of phenotypes that are not descendants of phenotype φ in the phenotype DAG. In a Bayesian Network, the local Markov assumption 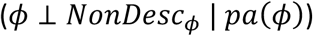 holds for every node φ. Intuitively, in the case of the HPO DAG, the conditional probability distribution associated with a phenotype φ defines the probability of observing φ given observations of its parent phenotypes *pa*(*φ*).

We define the conditional probability of observing phenotype *φ*, given its immediate parent(s), *pa(φ)*, as the number of genes associated with, *φ*,|*G*_*φ*_, over the number of genes associated with any 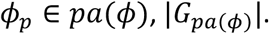 This follows from an assumption that each gene has equal probability of being mutated and resulting in the phenotype. Thus, the conditional probability distribution associated with each phenotype φ in the HPO DAG is defined as follows:

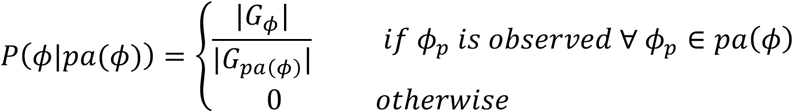

To compute the probability of any set of phenotypes Φ, following the chain rule of probability of events,

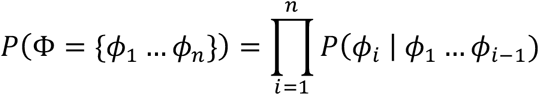

Given a set of phenotypes Φ including all ancestors (such that Φ = anc(Φ)), and using the local Markov assumption, this formula can be simplified to:

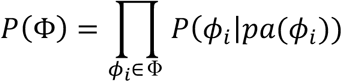

Substituting each factor in this multiplication with the appropriate element from the conditional probability distribution associated with each phenotype φ:

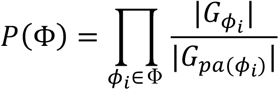

The information content (in bits) of a set of phenotypes φ is, by definition,

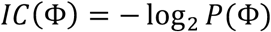

The Phrank score is defined as the information content of an intersection of two sets of phenotypes ancestral to the two input phenotype sets. One can use the above observations to derive an explicit formula to compute this quantity:

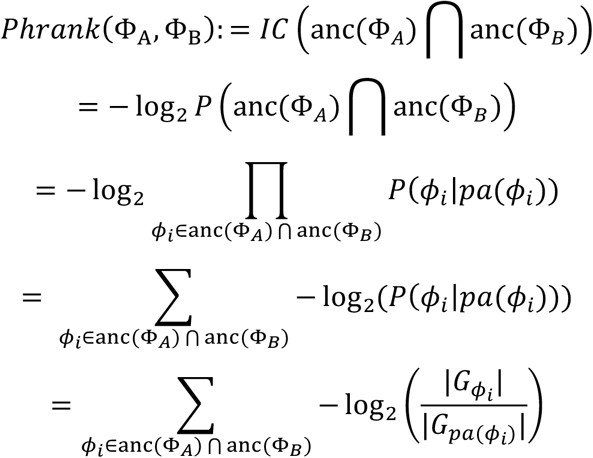

